# Resolving platypus behaviour from accelerometry: frequency-domain features improve detection of rhythmic behaviours in hydrodynamically challenging aquatic environments

**DOI:** 10.64898/2026.08.05.743151

**Authors:** Breony Webb, Michelle Ryan, Jessica L. Thomas

## Abstract

Developing robust methods to quantify how animals allocate time across behaviours is essential for understanding energy use, habitat requirements, and responses to environmental change. For cryptic, semi-aquatic mammals such as the platypus, direct observation is difficult, creating a reliance on remote biologging approaches that can reliably infer behaviour in the wild. However, aquatic environments can both smooth acceleration signals through hydrodynamic damping and introduce noise from water movement, turbulence, and drag, potentially obscuring behavioural differences of similar magnitudes.

We tested whether progressively incorporating biomechanical and frequency-domain (FFT-derived) predictors improved behavioural classification in hydrodynamically challenging aquatic environments. Tri-axial accelerometers were deployed on four ex situ platypuses, with synchronised video observations used to validate behaviour. From the acceleration data, we derived three predictor classes of increasing complexity: summary statistics describing activity level, engineered biomechanical variables capturing posture and body orientation, and FFT-derived features describing movement rhythm. These predictors were progressively incorporated into Random Forest models to classify five behaviours: burrow resting, surface resting, grooming, travelling/foraging, and diving.

Model performance improved with increasing predictor complexity, although gains were behaviour specific. FFT-derived features substantially improved classification of rhythmic behaviours such as diving and foraging, while engineered biomechanical predictors improved grooming detection. In contrast, resting behaviours, particularly surface resting, showed little improvement. Overall accuracy increased from ∼75% to ∼88% when frequency-domain features were included. Misclassification was greatest among behaviours with overlapping or low-amplitude signals, and cross-individual validation revealed reduced model generalisability, indicating that individual variation in movement patterns constrained transferability.

Incorporating frequency-domain features substantially improved behavioural classification in platypuses, particularly for rhythmic behaviours such as diving and foraging. This study provides the first validated accelerometry-based behavioural classification framework for the species and highlights the importance of matching predictor selection to behavioural mechanics. More broadly, the approach offers a transferable framework for aquatic and semi-aquatic taxa.

## 1. Introduction

Observing what animals do in their natural environment and how they allocate their time to each activity is central to ecology, yet it remains one of the greatest practical challenges in studying wildlife behaviour [1, 2]. Species that are difficult to observe directly are often nocturnal, wide-ranging, or inhabit environments where visibility is limited. These challenges are especially acute for semi-aquatic mammals, whose behaviour often occurs underwater or at night, beyond the reach of traditional field observation and ethogram-based approaches [3–5]. Understanding behaviour time allocation is particularly important for conservation because it links habitat use to energetic expenditure and survival [6]. For example, knowing how long animals spend foraging versus resting can indicate food availability, habitat quality, or disturbance effects [7]. In the case of some species these can be insights that cannot be obtained from location biotelemetry alone [8].

Biologging technologies have transformed this problem by enabling researchers to measure animal movement remotely [9, 10]. Among these tools, tri-axial accelerometers, small devices that are readily attachable to an animal, record acceleration along three spatial axes. The devices have become widely used because they capture the fine-scale body movements that underlie behaviour [11, 12]. By quantifying changes in posture [13, 12], motion size [14–16] and locomotor dynamics, behaviours such as resting, walking, swimming, diving, grooming and foraging have been elucidated, capabilities often unattainable from direct observation [3, 14, 12]. Consequently, accelerometry-based behavioural classification and modelling has been widely applied to birds [e.g. 12], mammals [e.g. 13, 17, 18], and aquatic taxa [e.g. 19–21], supporting studies of energetics [e.g. 8, 22], movement ecology and conservation applications [23].

While accelerometry provides a powerful approach for inferring behaviour in species where direct observation is limited, translating raw acceleration signals into meaningful behavioural categories requires analytical models that link sensor output to biomechanics [5, 12, 29]. Supervised machine-learning approaches, particularly Random Forests, support vector machines, and hidden Markov models, which learn relationships between acceleration-derived predictor variables and independently observed or video-validated behaviours [30–34]. Raw tri-axial acceleration data are typically summarised into predictor variables that capture movement magnitude (e.g. ODBA), body orientation, and temporal organisation [13–15, 35]. These variables allow models to distinguish behaviours not only by how much an animal moves, but by how movement is structured in space and time.

Predictor variables derived from simple summary statistics (e.g. mean and standard deviation) are effective for distinguishing active from inactive states but often perform poorly when behaviours share similar movement magnitudes [12]. To address this, studies increasingly incorporate biomechanical and frequency-domain variables that better represent how animals move [35–38]. Biomechanical variables describe how movement is distributed across body axes and capture posture and coordination [39, 40], while frequency-domain variables quantify the temporal structure, rhythm, and regularity of repeated movements [35, 36]. Frequency-domain characteristics underpin many aquatic behaviours, with movement often expressed as rhythmic oscillations such as stroke cycles, tail beats, or diving patterns [35, 39]. These frequency signals have been occasionally adopted to infer behaviour in marine and semi-aquatic taxa, including fish, seabirds, and diving mammals, where frequency metrics such as stroke rate are directly linked to locomotion and energetic expenditure [41, 42]. Despite this, the explicit incorporation of frequency-domain features derived via fast Fourier transforms (FFT) into machine learning classification frameworks remains limited, and their independent contribution relative to summary statistics and biomechanical predictors is rarely evaluated [43, 44]. This gap is particularly pronounced in semi-aquatic mammals, where hydrodynamic damping, introduced water noise, and continuous behavioural transitions can obscure acceleration magnitude, potentially elevating the importance of frequency-based descriptors where rhythmic behaviours (e.g. paddling, grooming, diving) may be more readily distinguished by their temporal structure than by movement magnitude alone.

Before applying such approaches to wild populations, it is necessary to validate the relationship between acceleration signals and behaviour under controlled conditions [12]. Ex situ individuals provide a critical opportunity to develop and test classification frameworks, and to assess their generalisability beyond the individuals on which they were trained [38, 45]. Generalisability remains a key challenge in accelerometry-based behavioural modelling, as classifiers often perform well within individuals but degrade across individuals or new datasets [45]. This reflects variation in morphology [46], movement style [12], sensor placement [29, 47], and model overfitting [45]. In semi-aquatic mammals, these challenges may be further compounded by environmental effects on signal transmission, highlighting the need to understand how individual-level biomechanics and signal structure interact to constrain model transferability.

The platypus (*Ornithorhynchus anatinus*) a semi-aquatic mammal presents an extreme case of observational constraint [24, 25]. It is largely nocturnal and spends most of its active time underwater foraging, surfacing only briefly (typically 10–15 s) to breathe and process prey before diving again [26]. Foraging is achieved by sweeping the bill laterally through benthic substrates while relying on electro- and mechano-sensory cues in their bill to detect prey [27]. Locomotion is highly specialised; propulsion during diving, swimming, and foraging is generated primarily by alternating forelimb strokes using large, webbed feet, whereas the hind limbs function mainly in steering and manoeuvring [24, 28]. This movement style is supported by a distinctive musculoskeletal anatomy, including a reptilian-like pectoral and pelvic girdle configuration with sprawling limb posture and robust girdle elements, which differs markedly from the parasagittal limb mechanics of most mammals [24, 28]. This structure facilitates powerful, multi-planar paddling in water [28]. The combination of aquatic foraging, brief surfacing events, and nocturnal activity means that most existing knowledge derives from radio-tracking and physiological studies describing coarse activity patterns or dive cycles [25, 26]. Consequently, fine-scale behavioural structure and time allocation among behaviours such as resting, grooming, foraging, and diving remain poorly understood.

To address the challenge that hydrodynamic damping and behavioural continuity obscure acceleration magnitude in semi-aquatic systems, we explicitly quantify the contribution of FFT-derived predictors within a unified modelling framework, using the platypus as a model system. There are currently no validated frameworks for accelerometry-based behavioural classification in the platypus. Here, we develop and validate a classification framework using high-resolution tri-axial accelerometry and synchronised video observations from *ex situ* individuals. Specifically, we evaluate the relative performance of predictor classes of increasing complexity; summary statistics, biomechanical variables, and frequency-domain (FFT-derived) features, to determine how each contributes to classification accuracy and generalisability across individuals. We test the hypothesis that frequency-domain features will improve the classification of behaviours characterised by rhythmic or cyclic movement (e.g. grooming, diving) but provide limited benefit for static or low-dynamic behaviours (e.g. surface resting). More broadly, this study establishes a transferable framework for behavioural classification in semi-aquatic systems and demonstrates how incorporating temporal structure can improve inference in environments and species where traditional time-domain metrics are constrained, providing a foundation for scaling biologging approaches to free-ranging populations.

## 2. Methods

### 2.1 Study animals and accelerometer deployment

Tri-axial accelerometer loggers were deployed on four captive platypuses (*Ornithorhynchus anatinus*; one male, three females) housed at Healesville Sanctuary, Victoria, Australia (Tab. 1). Loggers (MetaMotionS, MbientLab; 27 × 27 × 10 mm; 14 g) were dorsally attached to the rump under anaesthesia following established protocols [48]. Briefly, fur was trimmed at the attachment site, and devices were affixed to the skin using cyanoacrylate adhesive (Selleys Supa Glue), with surrounding fur repositioned to minimise hydrodynamic drag and tag-induced movement [11, 29]. Devices were aligned to record acceleration along the surge (anterior–posterior), sway (lateral), and heave (dorso–ventral) axes and calibrated following [29] (Fig. 1). Accelerometers recorded raw acceleration (±8 g) at 50 Hz and stored data onboard, with daily remote downloads via Bluetooth (BaseWare) during burrow rest, enabling non-invasive data retrieval.

**Figure 1.**
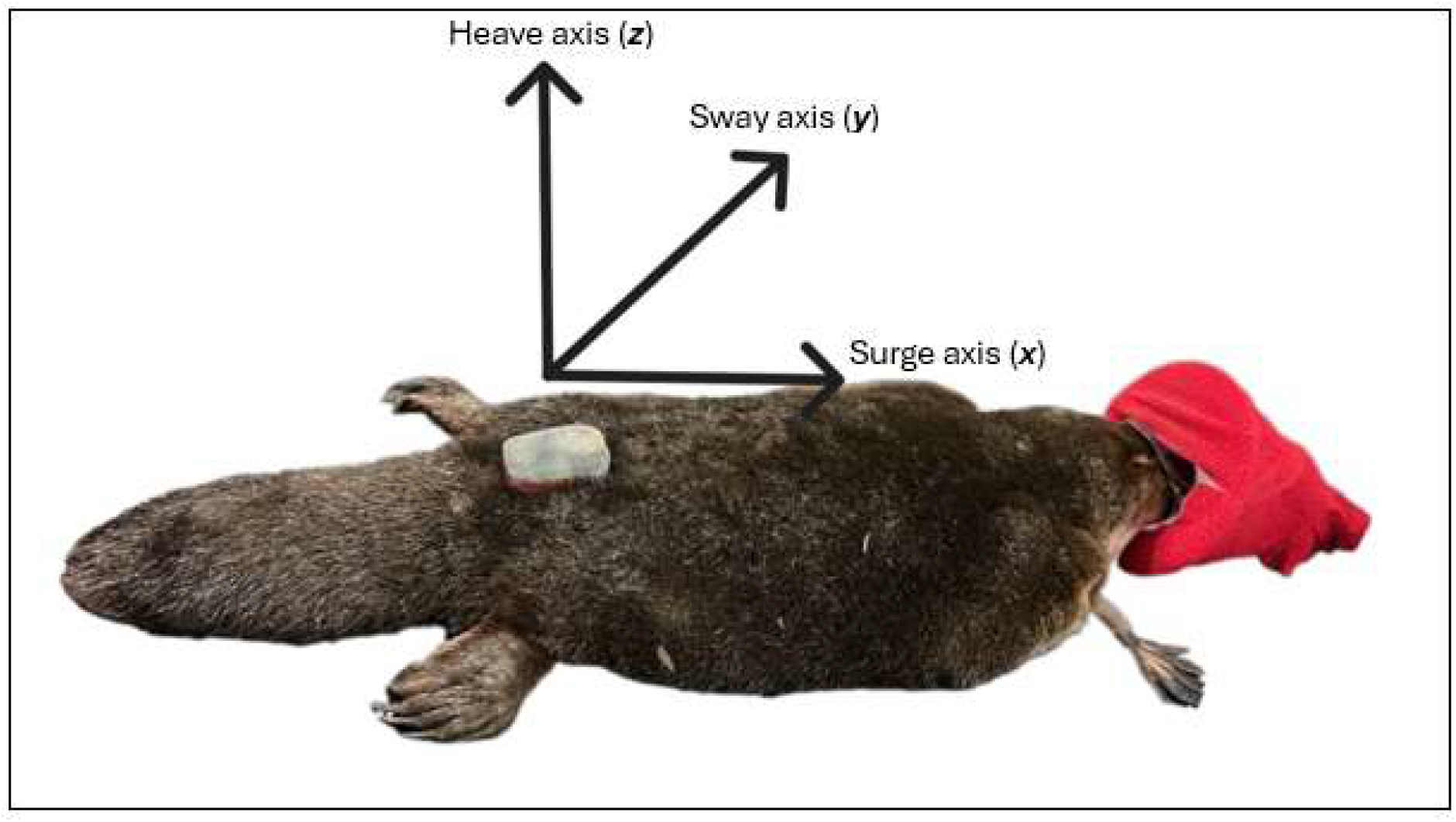
Tri-axial accelerometer placement on a platypus. Device attached dorsally on the rump, with axes aligned to surge (anterior–posterior), sway (lateral), and heave (dorso–ventral).

**Table 1.** Summary of study individuals, including sex, age, body mass, origin, and rearing history.

| Platypus ID | Sex | Age | Weight (g) | Birth type | Raised |
| --- | --- | --- | --- | --- | --- |
| A | Female | 12 Years 10 Months | 1222 | Captive | Parent |
| B | Female | 21 Years 11 Months | 1215 | Captive | Parent |
| C | Male | 21 Years 10 Months | 1635 | Wild | Hand reared |
| D | Female | 4 Years 10 Months | 1189 | Captive | Parent |

### 2.2 Behavioural observations and annotation

Behavioural observations were conducted over a two-week period using fixed in-house video systems (Swann Ultra HD DVR; 1–2 cameras per animal) within platypus enclosures. Video and accelerometer clocks were synchronised to align observed behaviours with acceleration data. Five behavioural classes were annotated: burrow rest, surface rest, grooming, foraging, and diving. Due to overlap in acceleration signals, surface travelling and foraging were combined into a single foraging category for model development. For each individual (N = 4), 10–12 video-verified examples per behaviour were extracted. Window lengths were tailored to behavioural duration: grooming, surface rest, and foraging (10–20 s), diving (1–3 s), and burrow rest (30–60 s).

### 2.3 Accelerometry analysis

#### 2.3.1 Summary statistic predictors

Static acceleration was calculated using a 1 s rolling mean [11, 15] and subtracted from raw acceleration to derive dynamic acceleration [16]. For each window, summary statistics describing movement intensity and axis-specific variability were calculated, including the mean and standard deviation of dynamic acceleration along the surge (X), sway (Y), and heave (Z) axes, as well as overall dynamic body acceleration (ODBA). Overall dynamic body acceleration was used as a proxy for movement and activity [13, 14] and calculated as the sum of absolute dynamic acceleration across axes [14]. Detailed formulae are provided in Supplementary Material S1.

#### 2.3.2 Engineered biomechanical and frequency-domain predictors

A suite of engineered predictors describing overall movement magnitude, body orientation, and relative within-window variability, following established biomechanical accelerometry frameworks (Tab. 2) [11, 12]. Predictors were calculated at the window level using combinations of axis-specific means and standard deviations of dynamic acceleration, capturing multi-axis organisation of movement to improve behavioural discrimination [16, 39]. Mathematical definitions are provided in Supplementary Material S2.

**Table 2.**
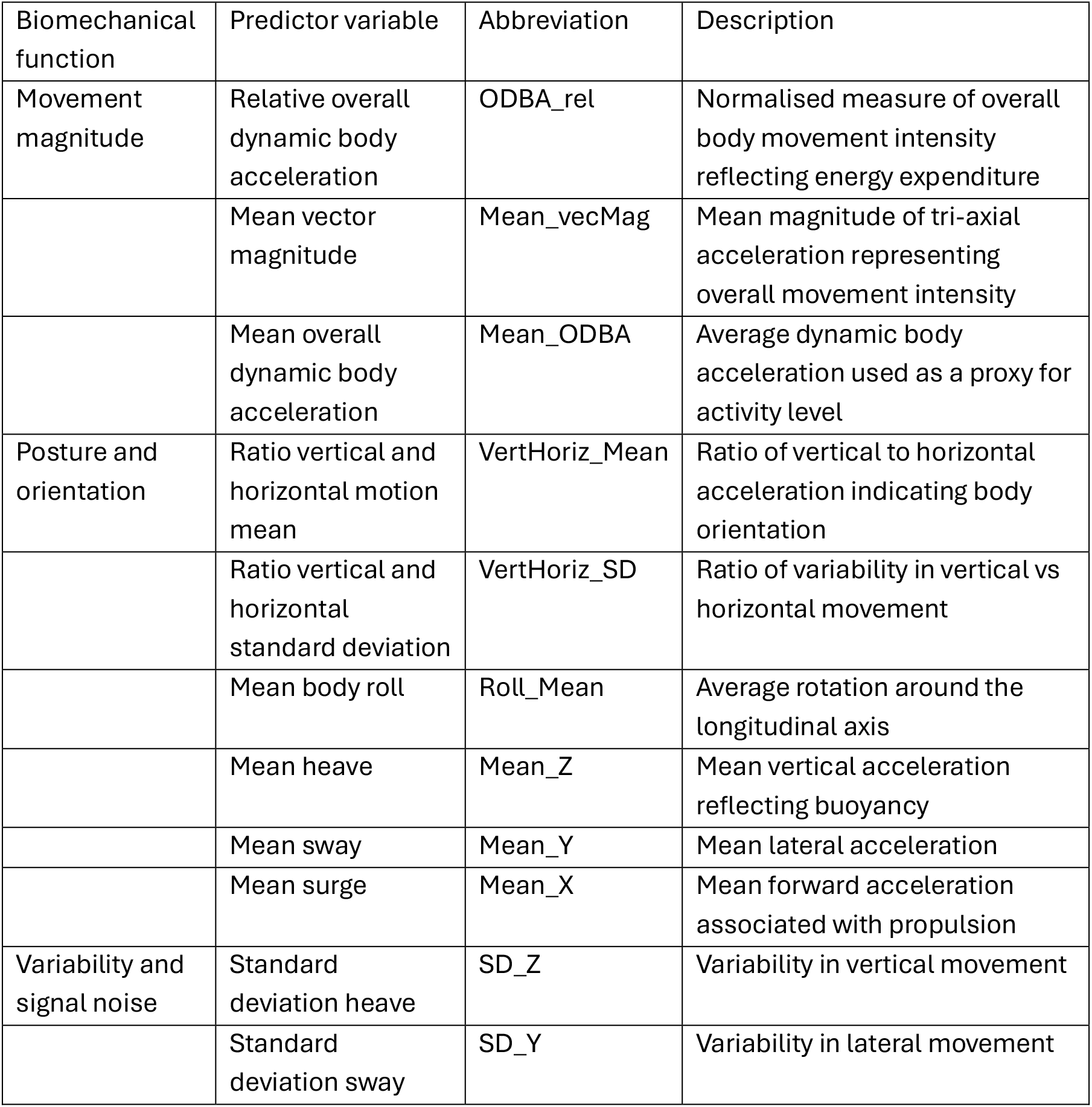
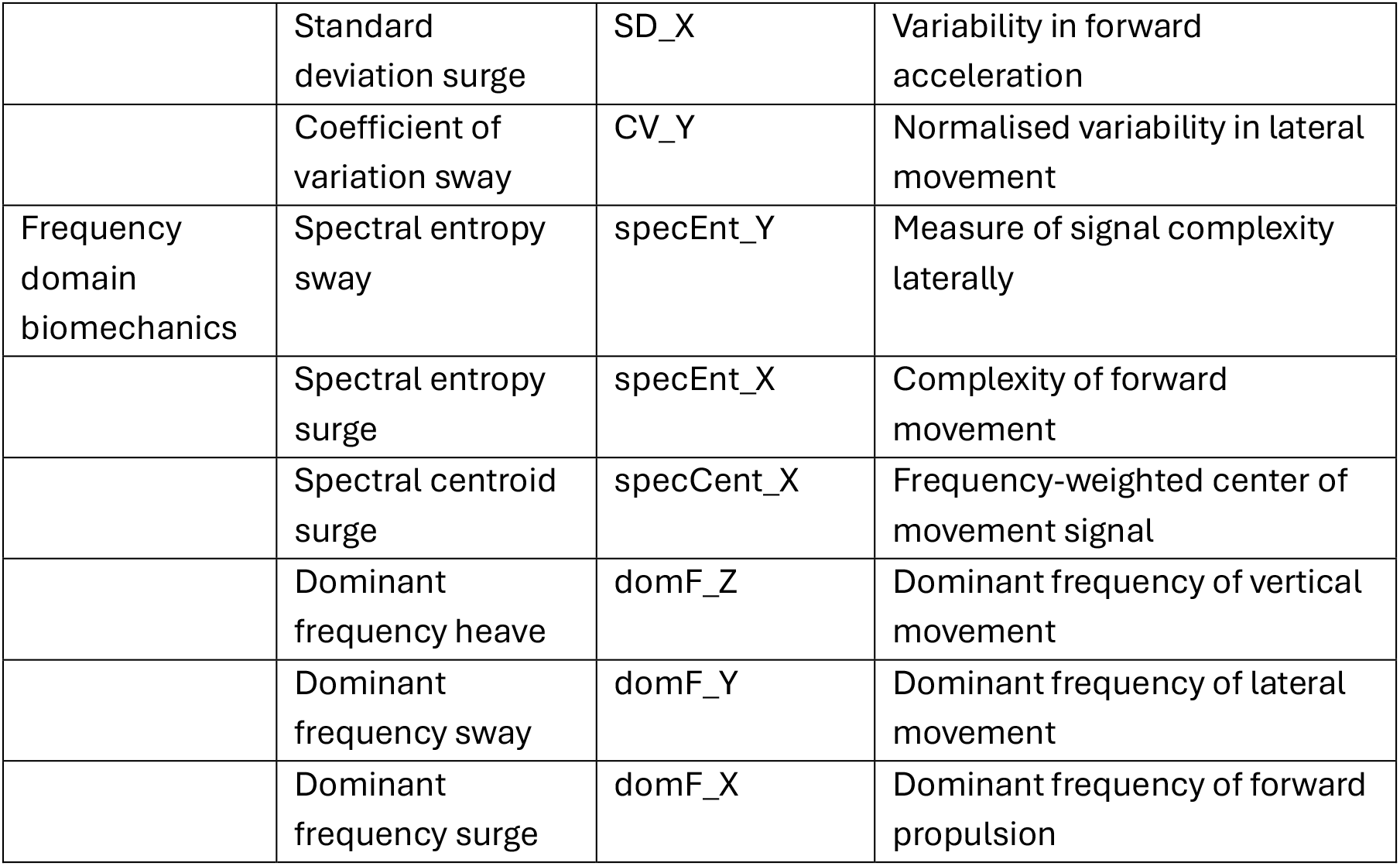
Accelerometry-derived predictor variables and biomechanical interpretation.

To characterise temporal structure beyond time-domain predictors, acceleration signals were transformed into the frequency domain using the Fast Fourier Transform (FFT) (Tab. 2) [35]. For each axis, dominant frequency, spectral centroid, and spectral entropy were extracted to quantify rhythmicity, tempo, and signal complexity, which are particularly informative for cyclic or irregular behaviours. Mathematical definitions are provided in Supplementary Material S3.

#### 2.3.3 Behavioural classification models and model validation

We trained three Random Forest (RF) classifiers differing in predictor complexity: (1) summary statistics only; (2) summary statistics plus engineered biomechanical predictors; and (3) summary, biomechanical, and FFT-derived frequency-domain predictors. This stepwise structure was used to isolate the contribution of increasingly complex, biologically informed predictors to classification performance. In preliminary analyses, individual predictor groups were also evaluated independently to assess standalone performance and redundancy, informing final model selection; combined models were retained as they improved classification accuracy. Random Forests were selected for their robustness to correlated predictors, non-linear relationships, and complex behavioural signatures [12, 30, 31]. To minimise multicollinearity, predictors were screened using correlation-based filtering and biological interpretability criteria.

All analyses were conducted in R (v4.5.1) [49], with data processing in dplyr and tidyr [50]. Models were implemented using the randomForest package [51] with consistent tuning parameters (2,500–3,000 trees; mtry = 3–4). Predictor importance was quantified using permutation-based scores and visualised as scaled heat maps (tidyverse; viridis), grouped by biomechanical function (Tab. 2).

Model performance was evaluated using a 70/30 train–test split and Leave-One-Subject-Out (LOSO) cross-validation to assess cross-individual generalisability [52]. Metrics included accuracy, sensitivity, specificity, Cohen’s κ, and out-of-bag (OOB) error rates [53, 54]. Behaviour-specific performance and predictor importance were calculated for each model.

#### 2.3.4 Outlier assessment and misclassification diagnostics

To assess whether misclassifications were driven by aberrant observations, we calculated an outlier score for each window as the maximum absolute z-score across all predictor variables [55]. Outlier score distributions were compared between correctly classified and misclassified windows using summary statistics and density plots (ggplot2; [56]). This allowed us to determine whether errors were associated with extreme signal values (e.g. tag placement artefacts) or reflected genuine behavioural overlap and within-behaviour heterogeneity [12, 29, 57].

#### 2.3.5 Multivariate structure and dispersion analyses

To examine the biomechanical structure underlying classification performance, we conducted multivariate ordination and dispersion analyses using standardised predictor variables. Principal Coordinates Analysis (PCoA; Euclidean distance) was used to visualise separation among behaviours and individuals in multivariate space [58–59]. Differences in within-group variability were assessed using permutational analysis of multivariate dispersion (betadisper), with permutation ANOVA and Tukey post-hoc tests to identify pairwise differences [60]. Analyses were performed using the vegan package [61], and figures generated with ggplot2 [56].

## 3. Results

### 3.1 Model performance

Five platypus behaviours (burrow rest, grooming, diving, surface resting, and foraging) were classified using acceleration signals and their derivatives. Random Forest performance improved with increasing predictor complexity (Tab. 3). Models using summary statistics alone achieved 75.36% accuracy (κ = 68.2%), increasing to 81.16% (κ = 75.61%) with engineered predictors. The full model, incorporating FFT-derived frequency variables, performed best, with the lowest OOB error (31.74%), highest accuracy (88.41%; κ = 86.89%), and strongest cross-individual generalisability (LOSO mean accuracy = 62.21%).

**Table 3.** Performance of three accelerometry classification models; (1) summary, (2) summary + engineered, and (3) summary + engineered + FFT. Evaluated using a 70/30 split and LOSO cross-validation.

| Validation statistics |  | 70/30 |  |  |  |  | LOSO |  |
| --- | --- | --- | --- | --- | --- | --- | --- | --- |
| Predictors | Model | Out-of-bag error rate | Accuracy | Specificity | Sensitivity | Cohen's Kappa | Accuracy | Cohen's Kappa |
| SD_Z<br>SD_X<br>SD_Y<br>Mean_ODBA<br>Mean_X<br>Mean_Z<br>Mean_Y | Summary statistics model only | 43.11 | 75.36 | 93.61 | 75.55 | 68.2 | 53.40 | 39.67 |
| CV_Y<br>SD_X<br>SD_Z<br>Mean_X<br>Mean_Z<br>Mean_ODBA<br>Mean_vecMag<br>Mean_Y<br>ODBA_rel<br>Roll_Mean | Engineered<br>+ summary<br>statistics<br>model | 34.73 | 81.16 | 94.92 | 82.39 | 75.61 | 55.10 | 41.15 |
| SD_Z<br>domF_Y<br>SD_X<br>SD_Y<br>Mean_ODBA<br>Mean_vecMag<br>domF_X<br>domF_Z<br>VertHoriz_SD<br>specEnt_X<br>Mean_Z<br>specCent_X<br>Mean_X<br>specEnt_Y<br>Mean_Y<br>VertHoriz_mean<br>CV_Y | FFT-derived +<br>engineered<br>+ summary<br>statistics<br>model | 31.74 | 88.41 | 96.75 | 86.89 | 84.79 | 62.21 | 50.76 |

### 3.2 Behaviour-specific classification performance and validation

Behavioural classification improved with increasing predictor complexity, with reduced confusion among behaviours (Tab. 4; Fig. 2). Summary statistic model achieved moderate to high accuracy (0.78–0.92), performing best for low-movement behaviours such as burrow rest (0.92) and surface rest (0.87). Incorporating engineered predictors improved accuracy for grooming (0.77–0.89) and diving (0.88–0.92), primarily through increased specificity (grooming: 0.67–0.95). The addition of FFT-derived predictors produced the largest gains for high-movement, rhythmic behaviours, increasing foraging accuracy by 10.4% (0.82–0.92) and diving by 9.9% (0.88–0.98), with corresponding improvements in sensitivity (foraging: 0.75–0.96; diving: 0.80–1.00). Burrow rest was consistently classified with near-perfect accuracy (0.92–1.00). Surface rest remained difficult to detect and decreased in accuracy as model complexity increased, with low sensitivity (∼0.64) despite increased specificity (0.93–1.00).

**Figure 2.**
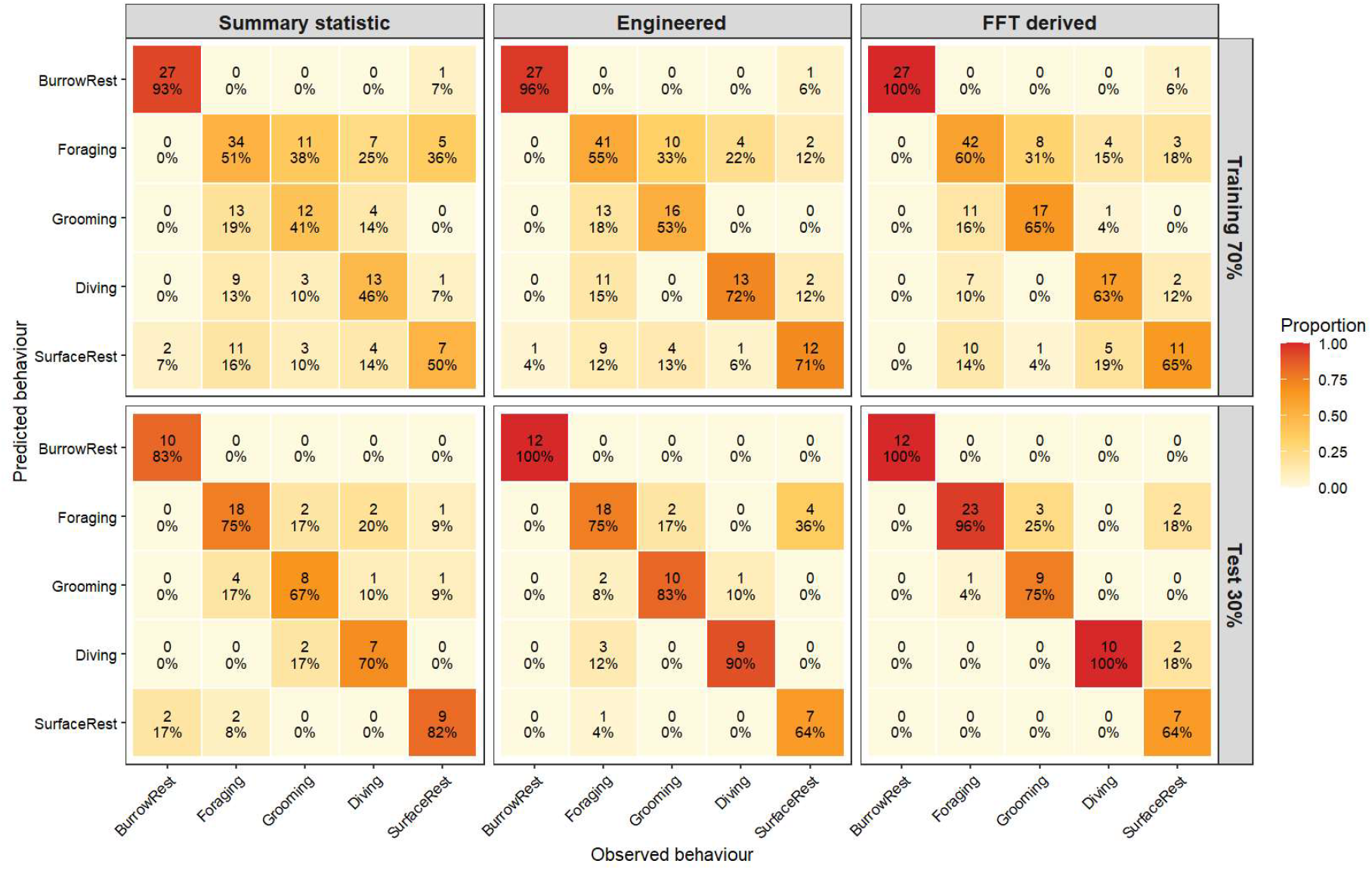
Confusion matrices for three accelerometry models (70/30 split). Columns show summary, engineered, and FFT models; rows show training (top) and test (bottom) data. Rows = predicted, columns = observed. Cells show counts (top) and row-wise % (bottom), with colour indicating class proportion.

**Table 4.** Behaviour-wise classification performance across model architectures.

| Model predictor variables | Behaviour | Accuracy | Sensitivity | Specificity |
| --- | --- | --- | --- | --- |
| Summary statistics | Burrow rest | 0.92 | 0.83 | 1 |
|  | Foraging | 0.82 | 0.75 | 0.89 |
|  | Grooming | 0.78 | 0.66 | 0.89 |
|  | Diving | 0.83 | 0.70 | 0.97 |
|  | Surface rest | 0.87 | 0.82 | 0.93 |
| Engineered + summary statistics | Burrow rest | 1 | 1 | 1 |
|  | Foraging | 0.80 | 0.75 | 0.87 |
|  | Grooming | 0.89 | 0.83 | 0.95 |
|  | Diving | 0.92 | 0.9 | 0.95 |
|  | Surface rest | 0.81 | 0.64 | 0.98 |
| FFT-derived + engineered + summary statistics | Burrow rest | 1 | 1 | 1 |
|  | Foraging | 0.92 | 0.96 | 0.89 |
|  | Grooming | 0.87 | 0.75 | 0.98 |
|  | Diving | 0.98 | 1 | 0.97 |
|  | Surface rest | 0.82 | 0.64 | 1 |

Overall, the addition of FFT predictors most improved detection of active behaviours characterised by repetitive, high-frequency motion (foraging, diving), while engineered predictors enhanced discrimination of posture- and variability-driven behaviours (grooming), and summary statistics performed best for low-movement states (surface rest). Misclassification patterns across training and test datasets suggest errors reflect genuine overlap in behavioural signatures rather than model bias, with lower test accuracy (training > test) likely due to sampling variability and class representation rather than overfitting (Fig. 2).

### 3.3 Predictor contributions across models

Predictor importance showed clear biomechanical structuring of accelerometry signals across models (Fig. 3), with behaviours characterised by distinct predictor families. In the summary-statistics model, behaviours were primarily discriminated by dynamic acceleration metrics, with SD_X, SD_Y, SD_Z, and Mean_ODBA dominating. Diving showed the highest dependence on Mean_ODBA and axis-specific SDs, consistent with powerful forelimb strokes and large heave/surge oscillations during propulsion.

**Figure 3.**
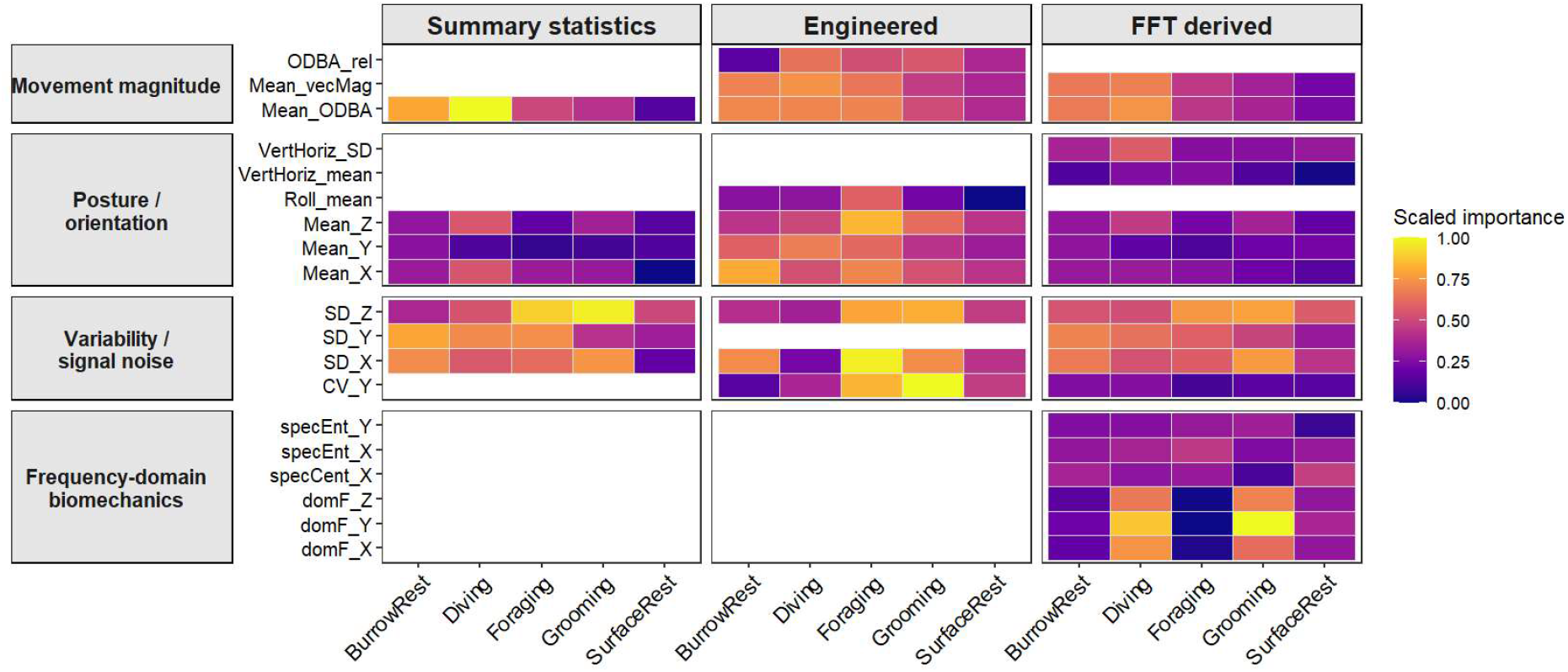
Behaviour-specific predictor importance across models. Heat maps of scaled variable importance for summary, engineered, and FFT models, grouped by biomechanical function.

With the inclusion of engineered predictors, importance shifted toward metrics capturing posture, movement magnitude, and variability (e.g. Mean_vecMag, Roll_mean, CV_Y). Grooming and foraging were better explained by fine-scale variability and rotational posture, reflecting irregular, multi-planar movements, while diving remained associated with high movement magnitude. Predictor contributions were more evenly distributed across behaviours in this model than simple acceleration summaries (Fig.3).

The FFT addition model produced the clearest behavioural separation, with frequency-domain predictors (e.g. domF_X, domF_Y, domF_Z) consistently ranking among the most influential for grooming and diving. Grooming was characterised by strong contributions from high-frequency components, reflecting rapid, oscillatory hindlimb movements, whereas diving exhibited moderate dominant frequencies coupled with high movement magnitude, consistent with stroke-driven propulsion. Foraging was associated with uniformly low-frequency signals across axes, indicative of slower, less periodic movement patterns. Surface rest occupied an intermediate position, with moderate frequency content and mixed contributions from entropy and centroid metrics, but overall low predictor importance, reflecting limited and less structured movement. (Fig.3).

### 3.4 Multivariate dispersion among behaviours

Multivariate dispersion differed significantly among behaviours (betadisper ANOVA: F(4, 231) = 23.06, p < 0.001), indicating variation in within-behaviour accelerometry signatures. Grooming and foraging showed the lowest dispersion, forming compact clusters in PCoA space and reflecting highly stereotyped movements (Fig. 4). In contrast, diving exhibited greater dispersion, consistent with variable locomotor dynamics linked to changes in depth, orientation, and propulsion. The highest dispersion occurred in burrow rest, and to a lesser extent surface rest, which formed diffuse and elongated clusters in multivariate space (Fig. 4).

**Figure 4.**
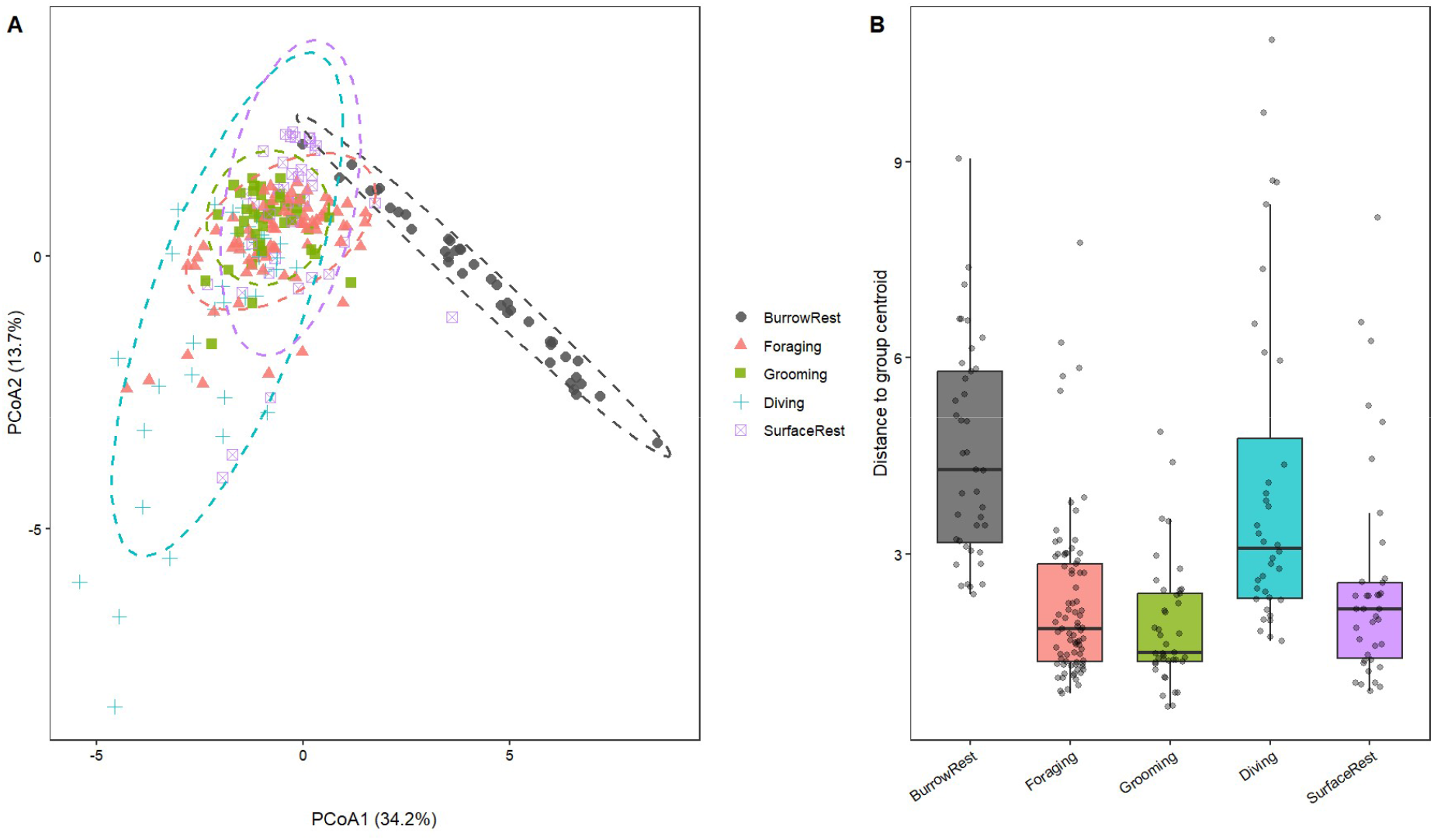
Multivariate dispersion across behaviours. (A) PCoA of standardised predictors with 95% ellipses showing within-behaviour variability. (B) Distances to behavioural centroids (betadisper ANOVA: F(4, 231) = 23.06, p < 0.001).

### 3.5 Outliers and misclassification

Misclassified windows had lower central tendency in lower outlier scores (median = 13.5, mean = 78.5) than the dataset overall (median = 61.1, mean = 180.8) and were not overrepresented in the upper tail of the distribution (Fig. 5). Extreme outliers (maximum |z| > 300–1600) were predominantly classified correctly. Density distributions overlapped substantially between correctly classified and misclassified windows, with no enrichment of extreme values among misclassifications (Fig. 5).

**Figure 5.**
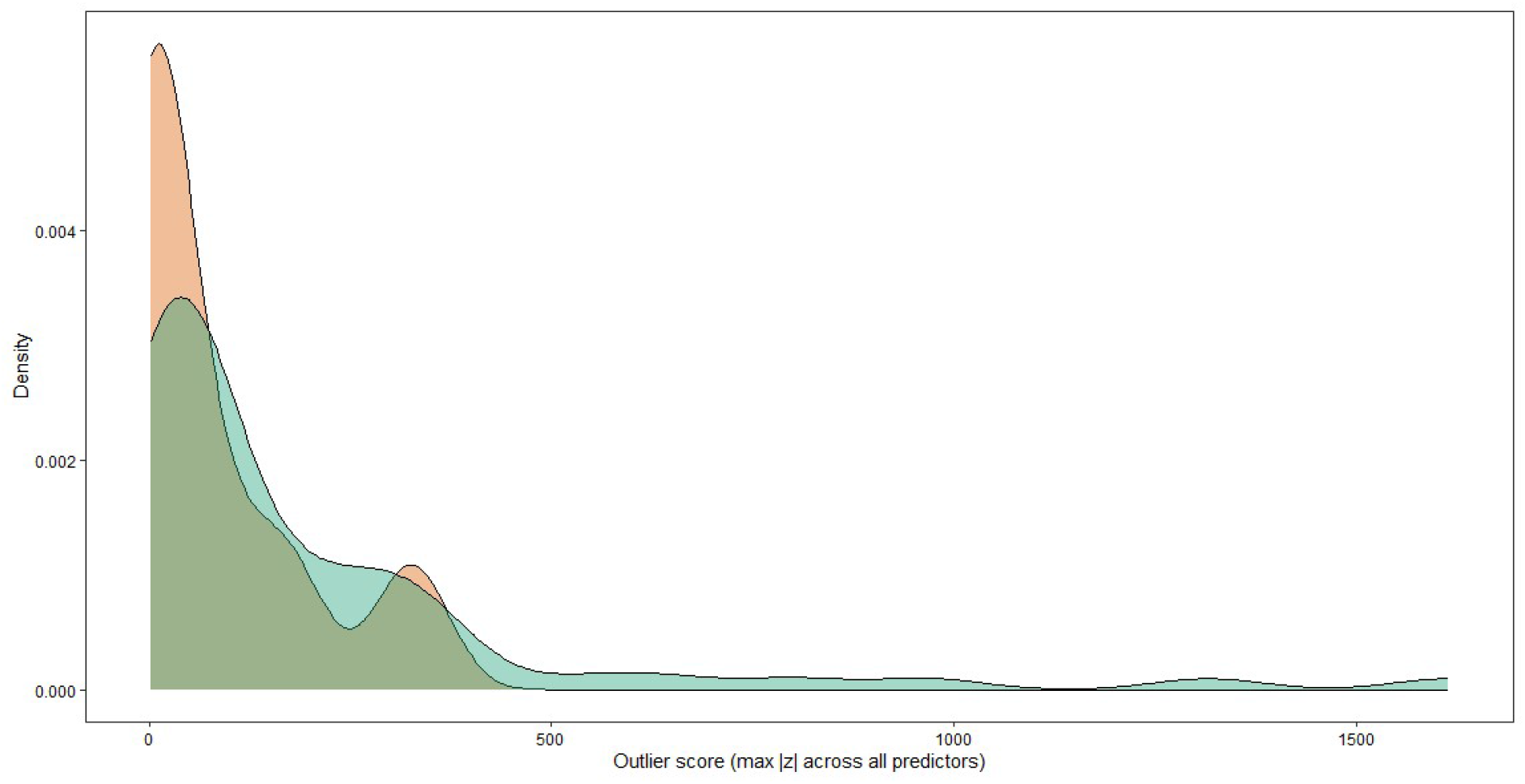
Density distributions of maximum absolute z-scores across all predictors. Misclassified (orange) windows do not exhibit elevated outlier scores relative to correctly classified windows (green), indicating errors reflect behavioural overlap rather than extreme segments.

### 3.6 Individual differences and model transferability

Individuals exhibited structured differences in accelerometry predictors and classification performance, with multivariate dispersion differing significantly among individuals (betadisper ANOVA: F (3, 85.18) = 5.38, p < 0.001), indicating systematic individual effects. In PCoA space, observations clustered by individual identity, with distinct centroid offsets and partially non-overlapping dispersion ellipses across individuals (Fig. 6). These patterns were consistent across behaviours, suggesting persistent individual differences within behavioural states.

**Figure 6.**
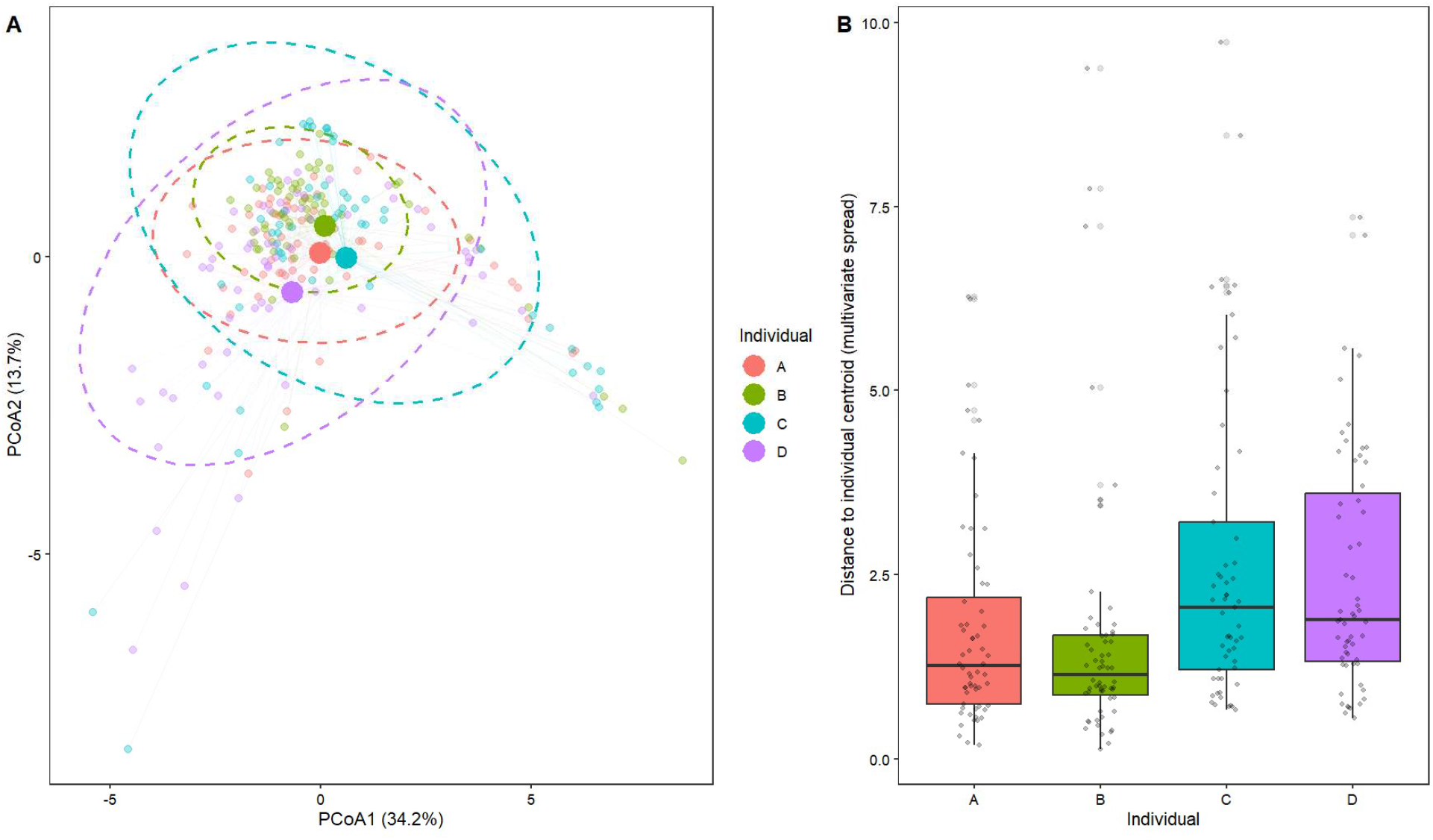
(A) Principal coordinates analysis (PCoA) of combined summary, engineered, and FFT-derived predictors, coloured by individual, with points representing behavioural windows and dashed ellipses showing 95% confidence regions. (B) Distances to individual centroids, summarising within-individual dispersion (betadisper ANOVA: F(3, 85.18) = 5.38, p < 0.01).

Individual-level offsets quantified across all predictors further supported this interpretation. Mean dynamic acceleration (ODBA), axis-specific means, and frequency-domain metrics differed consistently among individuals, with some individuals (e.g. Platypus C and D) exhibiting higher within-individual dispersion across multiple predictors, while others (e.g. Platypus A and B) showed more compact, stereotyped signal structure. These differences were not restricted to any single biomechanical domain, but spanned posture-related (mean X/Y/Z), variability-based (SDs, CVs), and frequency-domain descriptors, indicating individual-specific movement signatures.

These effects reduced model generalisability. LOSO validation of the FFT addition model showed lower performance relative to within-individual validation, with accuracy ranging from 0.56 (Platypus D) to 0.70 (Platypus B) (κ = 0.41–0.61, respectively). Individuals with higher dispersion and greater deviation from the population centroid (notably Platypus C and D) had the lowest accuracy (Fig. 6; Tab. 3), indicating reduced transferability of classifiers across individuals. Post hoc comparisons confirmed significant centroid differences involving Platypus D relative to Platypus A (Δ = 1.20, p = 0.024) and Platypus B (Δ = 1.51, p = 0.002), while other contrasts were weak or non-significant.

## 4. Discussion

This study presents the first accelerometry-based behavioural classification framework for the platypus, showing that biomechanically informed time- and frequency-domain predictors improve classification accuracy, interpretability, and cross-individual generalisability for active and rhythmic behaviours. Using tri-axial accelerometry and Random Forest models, we classified five behaviours; burrow resting, surface resting, grooming, foraging, and diving, across models of increasing complexity.

### 4.1 Comparative performance and biomechanical drivers of accuracy

Overall accuracy achieved by the FFT-augmented model (88% under random splits; ∼62% under LOSO) is comparable to, or exceeds, performance reported for many aquatic and semi-aquatic taxa using similar machine learning approaches. Studies of diving seabirds, marine mammals, fish, reptiles, and semi-aquatic mammals commonly report overall accuracies between ∼70–95% when classifying broad behavioural states [e.g. 18, 19, 44, 62-64]. Consistent with these systems, behaviours characterised by strong forelimb propulsion and rhythmic rutter like hindlimb movement close to logger body placement, particularly diving, were classified with the highest accuracy in our models, especially once frequency-domain predictors were included. Although, foraging was classified with relatively high accuracy, grooming and surface resting were often misclassified as foraging, which were also reported in accelerometry studies on fur seals [34, 47]. Similarly, burrow resting was consistently accurately identified, with minimal misclassification of other behaviours as burrow resting. This may be due to the relatively low-movement signature compared to all other behaviours [12] and the equal number/duration of training data applied across each behaviour [65]. Dunford et al., (2024) [66] notes that RF models trained on behaviourally imbalanced datasets often favour dominant behaviours, leading to poorer classification of rare behaviours. For example, on free ranging cattle misclassification of rare behaviours such as rest has occurred [67]. We kept the model to five behaviours classes commonly observed in platypuses, expanding the model to include rarer behaviours such as courtship and mating with subsequent imbalance data duration to train on, may see the misclassification of them into other more abundant behaviours.

In contrast, grooming and surface resting were more difficult to classify reliably. Grooming involves irregular patterns of repetitive forelimb and hindlimb strokes, and multi-planar movements expressed as lateral rolling and sway as the animal changes body position and limb used to groom, rather than clean, repetitive stroke cycles observed in foraging and diving. Similar challenges have been reported in beavers, where grooming exhibited high lateral dispersion due to alternating left to right body movements [19], and in terrestrial mammals and birds where body-maintenance behaviours showed weakly stereotyped acceleration signals [17, 68]. In platypuses, this challenge is exacerbated by the rump-mounted sensor that capture trunk motion and fine-scale hindlimb movement but attenuate fine-scale forelimb and cranial movements. As a result, forelimb water treading during surface resting, were often confused with low-amplitude swimming during foraging or subtle postural adjustments while swimming.

Diving in platypus is initiated with strong forelimb propulsion and head ducking into the water from a surface rest position. Diving produced highly structured acceleration signals driven by strong heave propulsion, rhythmic paddling and posture change, which were effectively captured by engineered and frequency-domain predictors. Misclassification of surface resting as diving likely reflects transitional windows overlapping dive initiation as their movement intensities contrast, highlighting a key constraint: behavioural states in accelerometry data are continuous rather than discrete [12, 19]. During surface rest, the animal may float motionless, paddle lightly to maintain position, adjust posture relative to surface conditions, or transition rapidly into travel or diving. In this study, these behaviours during surface rest generated weak, low-energy acceleration signals with substantial overlap among other behaviours states, inflating within-class dispersion and reducing model separability from other behaviours. Similar misclassification between resting type behaviours and low-movement behaviours [34], but also low-movement phases of diving ascents in sea turtles, where slow ascents were frequently confused with resting [63]. In our study, surface resting also varied markedly among individuals in both duration and mean ODBA, with some individuals rarely entered a prolonged “true” surface resting state.

Together, these results demonstrate that classification accuracy is not solely determined by model choice, but by how strongly behaviours are expressed in each biomechanical signal and successfully captured by the sensor position. Improving discrimination among grooming, foraging, and surface states will likely require complementary sensors (e.g. wet–dry sensors; [18]) or alternative tag placement targeting head, bill, or forelimb motion [20, 69], which underpin key functional behaviours in platypuses [24, 28].

### 4.2 Contribution of engineered and frequency-domain predictors

Models based solely on summary acceleration statistics perform well at classifying active from non-active behaviours but often struggle to discriminate behaviours with overlapping movement intensities or postures, particularly in animals with irregular and highly transitional states [5, 15, 35]. Our results in the platypus mirror this limitation. The addition of engineered predictors improved discrimination by capturing posture, relative movement magnitude, and variability, particularly enhancing specificity for behaviours prone to confusion such as grooming [5, 12, 40].

Further gains were achieved with FFT-derived predictors, which improved detection of rhythmic propulsion behaviours such as diving and grooming by capturing tempo-frequency structure. This aligns with previous work demonstrating the value of spectral features for identifying rhythmic and complex behaviours in aquatic systems [5, 35, 36]. Dominant frequency (domF) may have been more informative as a predictor for foraging if it were mounted closer to movement source (forelimbs and head movement) [59], as rump-mounted sensors will attenuate fine-scale oscillations from forelimbs and head movements [11, 15, 20, 59].

Notably, classification performance for surface resting declined with increasing model complexity due to reduced sensitivity despite improved specificity. This reflects a key trade-off: as models incorporate more structured biomechanical information, they become more conservative in assigning behaviours lacking consistent signal structure [70]. Surface resting, as a weakly defined and transitional state, therefore becomes increasingly difficult to detect. This highlights that increasing model complexity can expose intrinsic limits in behavioural separability rather than universally improving classification accuracy.

The use of Random Forests in this study reflects their suitability for capturing non-linear, multi-dimensional relationships between biomechanical predictors and behaviour, rather than model optimisation *per se.* While alternative machine learning approaches may yield comparable performance, the patterns observed here are driven primarily by the structure of the underlying biomechanical signal.

### 4.3 Biomechanical and individual-level constraints on generalisability

Misclassification errors in our models were driven primarily by genuine overlap among behavioural acceleration signatures rather than extreme or pathological outliers, consistent with findings across taxa [12, 34]. Behaviours with diffuse multivariate structure, short duration (e.g. surface rest for some individuals), transitional (e.g. foraging), or weakly stereotyped states (e.g. grooming), produced overlapping predictor distributions that inherently limited clear separability, irrespective of model complexity.

Model performance declined under LOSO validation, adding evidence to individual differences in movement biomechanics. Individuals with greater within-individual variability and displacement from the population centroid showed reduced classifier transferability (e.g. Platypus D), reinforcing that individual biomechanics impose fundamental limits on generalisation [70]. As outlined in the introduction, such variation is widely reported and arises from differences in morphology, movement execution [12], and sensor placement and data processing protocols [29, 66]. Given standardised deployment protocols and consistent sampling design, these differences are unlikely to reflect measurement bias and instead represent intrinsic variation in how behaviour is expressed biomechanically. These findings highlight a core challenge in accelerometry-based behavioural classification: models trained on one set of individuals may not fully generalise to others unless this variation is explicitly accounted for through larger training datasets, individual-normalised predictors, or hierarchical modelling approaches.

### 4.4 Implications and conclusions

Platypuses present exceptional challenges for behavioural ecology due to their aquatic and predominantly nocturnal lifestyle [25], where direct observation is limited and acceleration signals are shaped by hydrodynamic damping, overlapping movement magnitudes and continuous movement transitions. By explicitly comparing predictor classes, this study demonstrates that incorporating temporal structure through frequency-domain (FFT-derived) features substantially improves behavioural classification in this system, but in a behaviour-dependent manner. Improvements were strongest for dynamic behaviours characterised by rhythmic or cyclic movement, such as diving and foraging, while static or low-dynamic behaviours such as surface resting showed little or no improvement, performing best under basic summary statistics. This highlights that the value of frequency-domain features is contingent on the underlying biomechanics of behaviour, rather than universally enhancing classification performance making the case for hierarchical models [43]. At the same time, misclassification among behaviours with low-amplitude or overlapping signals indicates that limitations arise from intrinsic similarities in how behaviours are expressed and recorded, rather than deficiencies in model structure alone. Strong individual effects further demonstrate that variation in movement style and biomechanics constrains model generalisability, reinforcing the need to account for individual-level differences when applying classification models to new datasets or wild populations.

Finally, by isolating the contribution of frequency-domain features within a unified modelling framework, it provides a mechanistic basis for selecting predictor variables according to behavioural structure. This has wider implications for aquatic and semi-aquatic taxa, where incorporating temporal dynamics may be essential for resolving behaviours that are otherwise indistinguishable using traditional time-domain approaches, and for improving the ecological interpretation of biologging data in complex environments.

## Acknowledgements

We acknowledge the contribution of Healesville Sanctuary Veterinarian staff and Zookeepers, particularly Angelica Aguilar for assisting with animal housing set-up for the project. We acknowledge the statistical advice provided by Christopher Howden and minor manuscript feedback provided by Dr John Hunt - Head of Discipline, Life Sciences, School of Science Western Sydney University.

## Author contributions

B.W., J.T. and M.R. conceived the idea and designed the methodology. B.W. collected the data and conceptualised the modelling approach for analysis; prepared and analysed the data and wrote the code for the modelling and validation and led writing of the manuscript. All authors contributed critically to the drafts and gave final approval for publication.

## Data availability statement

The data associated with the study are available at reasonable request from corresponding author. Video recordings used for behavioural validation are not publicly available as per institutional policy but may be accessed upon reasonable request subject to institutional approval.

## Ethics approval

All experimental procedures were conducted in accordance with institutional and national guidelines for animal welfare and were approved by the Zoos Victoria Animal Ethics Committee (Approval No. ZV24016). The study was conducted on animals under the care of Zoos Victoria, and all necessary institutional permissions for their inclusion in research and publication were obtained.

