## Supplementary Materials for "Resolving platypus behaviour from accelerometry: frequency-domain features improve detection of rhythmic behaviours in hydrodynamically challenging aquatic environments"

**S1. Tri-axial accelerometry and data segmentation**

For each axis, summary statistics describing movement intensity and variability were calculated at the window level, including the mean and standard deviation of dynamic acceleration (Shepard et al., 2008; Wilson et al., 2006):

$$\mathrm{Mean}_{X}, \mathrm{Mean}_{Y}, \mathrm{Mean}_{X}$$

$$\mathrm{SD}_{X}, \mathrm{SD}_{Y}, \mathrm{SD}_{X}$$

Overall Dynamic Body Acceleration (ODBA) was calculated following established methods:

$$ODBA=\mid A_{X}\mid+\mid A_{Y}\mid+\mid A_{Z}\mid$$

where A_X_, A_Y_, A_Z_ are dynamic acceleration components along each axis (Wilson et al., 2006; Gleiss et al., 2011).

**S2. Engineered biomechanical predictors**

To capture biomechanically meaningful structure beyond axis-specific summary statistics, a suite of engineered predictors describing movement magnitude, body orientation, and relative within-window variability was derived from window-level summary metrics (Nathan et al. 2012).

$$Mean\_vecMag = \sqrt{\mathrm{Mean}_{X}^{2} + \mathrm{Mean}_{Y}^{2} + \mathrm{Mean}_{Z}^{2}}$$

and the Euclidean norm of the variability vector (Nathan et al., 2012):

$$SD\_vecMag = \sqrt{\mathrm{SD}_{X}^{2} + \mathrm{SD}_{Y}^{2} + \mathrm{SD}_{Z}^{2}}$$

A normalised dynamic acceleration metric was calculated as:

$$ODBA\_rel = \frac{Mean\_ODBA}{Mean\_vecMag + \varepsilon}$$

where ε= 10^-6^ prevents division by zero.

Body orientation was approximated using pitch- and roll-like proxies derived from mean acceleration components (Shepard et al., 2008):

$${Pitch}_{mean} ={tan}^{-1}\frac{{Mean}_{Z}}{\sqrt{{Mean}_{X}^{2}+{Mean}_{Y}^{2}}}$$

$${Roll}_{mean} ={tan}^{-1}\frac{{Mean}_{Y}}{\sqrt{{Mean}_{X}^{2}+{Mean}_{Z}^{2}}}$$

Relative within-window variability was quantified using coefficients of variation for each axis:

$$\mathrm{CV}_{X}=\frac{\mathrm{SD}_{X}}{\left| \mathrm{Mean}_{X} \right|+\varepsilon}$$

$$\mathrm{CV}_{Y}=\frac{\mathrm{SD}_{Y}}{\left| \mathrm{Mean}_{Y} \right|+\varepsilon}$$

$$\mathrm{CV}_{Z}=\frac{\mathrm{SD}_{Z}}{\left| \mathrm{Mean}_{Z} \right|+\varepsilon}$$

Vertical–horizontal contrasts in movement and posture were captured using:

$$VertHoriz\_SD=\frac{\mathrm{SD}_{Z}}{\mathrm{SD}_{X}+\mathrm{SD}_{Y}+\varepsilon}$$

$$VertHoriz\_Mean=\frac{\left| \mathrm{Mean}_{Z} \right|}{\left| \mathrm{Mean}_{X} \right|+\left| \mathrm{Mean}_{Y} \right|+\varepsilon}$$

**S3. Frequency-domain feature extraction**

To capture temporal structure not represented in time-domain features, acceleration signals along each axis were transformed into the frequency domain using the Fast Fourier Transform (FFT) (Cooley and Tukey 1965). For a window of (N) samples, the discrete Fourier transform was defined as:

$$X_{k}=\sum_{n=0}^{N-1} x_{n}e^{-2\pi ikn/N}$$

The power spectrum was computed as:

$$P_{k}=\left| X_{k} \right|^{2}$$

Only positive frequencies were retained:

$$f_{k}=\frac{kf_{s}}{N}, k=1,\ldots, \left\lfloor\frac{N}{2} \right\rfloor$$

All FFT features were calculated independently for each axis (X, Y, Z).

From the power spectrum, three descriptors were extracted for each axis (Nathan et al. 2012):

**Dominant frequency (domF)**

$$domF=f_{k^{*}}, k^{*}=\begin{matrix} argmax(P_{k}) \\ k \end{matrix}$$

**Spectral centroid (specCent)**

$$\mathrm{specCent}=\frac{\sum_{k} f_{k}P_{k}}{\sum_{k} P_{k}}$$

**Spectral entropy (specEnt)**
First, the spectrum was normalised:

$$p_{k}=\frac{P_{k}}{\sum_{k} P_{k}}$$

Shannon entropy was calculated as (Shannon 1948):

$$H=-\sum_{k} p_{k}\log(p_{k})$$

and normalised:

$$\mathrm{specEnt}= \frac{H}{\log K}$$

where (K) is the number of retained frequency bins, yielding:

$$0\leq\mathrm{specEnt}\leq1$$
